# Different extrapolation of moving object locations in perception, smooth pursuit and saccades

**DOI:** 10.1101/2022.10.26.513821

**Authors:** Matteo Lisi, Patrick Cavanagh

## Abstract

The ability to accurately perceive and track moving objects is crucial for many everyday activities. In this study, we use a “double-drift stimulus” Lisi and Cavanagh (2015); Shapiro et al. (2010); Tse and Hsieh (2006) to explore the processing of visual motion signals that underlie perception, pursuit, and saccade responses to a moving object. Participants were presented with peripheral moving apertures filled with noise that either drifted orthogonally to the aperture’s direction or had no net motion. Participants were asked to saccade to and track these targets with their gaze as soon as they appeared, and then to report their direction. In the trials with internal motion, the target disappeared at saccade onset so that the first 100 ms of the post-saccadic pursuit response was driven uniquely by peripheral information gathered before saccade onset. This provided independent measures of perceptual, pursuit, and saccadic responses to the double-drift stimulus on a trial-by-trial basis. Our analysis revealed systematic differences between saccadic responses on one hand and perceptual and pursuit responses on the other. These differences are unlikely to be caused by differences in the processing of motion signals because saccade and pursuit appear to use a common motion processing mechanism (e.g., Erkelens, 2006; Fleuriet and Goffart, 2012). We conclude that our results are instead due to a difference in how the processing mechanisms underlying perception, pursuit, and saccades combine motor signals with target position. These findings advance our understanding of the mechanisms underlying dissociation in visual processing between perception and eye movements.

## Introduction

Visual motion serves multiple functions in visual perception, including object tracking, foreground-background segmentation Born et al. (2000), depth perception through motion parallax Rogers and Graham (1979), and computation of 3D structure Jain and Zaidi (2011). Beyond perception, visual motion plays a critical role in motor control, such as smooth pursuit eye movements Lisberger (2010), online adjustments of hand movements Gomi (2008); Whitney et al. (2003), and posture control Kelly et al. (2005); Lee and Lishman (1975). Estimating the velocity of moving objects is crucial for accurate planning of actions directed toward object in motion. Even saccadic eye movements, the fastest goal-directed action that humans can produce, are informed by velocity information when directed toward a moving object Etchells et al. (2010); Gellman and Carl (1991); Groh et al. (1997).

When tracking moving objects the brain uses motion signals to correct for the displacement of targets and guide interceptive motor actions. Evidence suggests that similar corrective adjustments are implemented in perception, where moving objects are usually perceived ahead of their physical location in space De Valois and De Valois (1991); Ramachandran and Anstis (1990); Whitney (2002). However, whether perception and motor control are guided by the same visual processing mechanisms is a topic of debate Cardoso-Leite and Gorea (2010); Spering and Carrasco (2015).

In a previous study, we investigated the processes that determine the perceived location of a moving object and those guiding saccadic eye movements toward the same object Lisi and Cavanagh (2015). We found that these processes are different. Participants were asked to make eye movements and judge the location of a moving stimulus whose perceived and physical direction of motion were dissociated, producing an error in the predictions of its future locations. The stimulus was a moving aperture containing a drifting pattern that moved along a direction 90°apart from the physical direction of displacement of the aperture (a double-drift stimulus). While perceptual judgments revealed the accumulation of a location error along the perceived (as opposed to physical) direction of motion over a long interval Kwon et al. (2015), interceptive eye movements were much less influenced by the perceived motion direction and showed only a small, constant error that did not vary over time. We found that the perceived trajectory could be recovered from the distributions of saccade landing positions only when a delay was introduced between the presentation of the stimulus and the onset of the movement, so that saccadic responses were memory-guided rather than based on online visual input Massendari et al. (2018). This suggests that the visual information that allows the saccadic system to avoid the accumulation of position error must be short-lived, possibly related to the fast decay of neural activity observed in oculomotor structures after the disappearance of visual targets Edelman and Goldberg (2001). Indeed, we found that the dissociation with perceived location was limited to interceptive saccades. Hand pointing movements, even when the pointing latency was matched to that of eye movements, revealed a systematic bias toward the perceived trajectory Lisi and Cavanagh (2017b).

Saccades are not the only eye movements that differ from perception. Pursuit eye movements, in some cases, reveal motion estimates that deviate from perceptual judgments Simoncini et al. (2012); Spering et al. (2011). To compare motion estimates used in saccades, pursuit, and perception, we developed a novel experimental paradigm that allowed us to measure pursuit, perceptual, and saccadic responses to a peripheral double-drift stimulus simultaneously. Our results indicate that perceptual direction judgments and pursuit responses have a strong similarity, as they both display large deviations from the physical direction of the target. On the other hand, saccade landing positions show only a small bias, which suggests extrapolation along a direction closer to the physical direction of the target, or extrapolation along the perceived direction but for a shorter interval than the saccadic latency. Saccades and pursuit are usually interdependent and cooperate to track moving objects Erkelens (2006); Fleuriet and Goffart (2012); Goettker et al. (2018); Lisi and Cavanagh (2017a), so it is unlikely that they use different estimates of target velocity. Thus, our results suggest that the dissociations between saccades and perceptual judgments in the localization of moving objects Lisi and Cavanagh (2015) are not due to differences in the processing of motion signals but in how velocity information is used to extrapolate future target positions.

## Material and Methods

### Participants

Eight subjects participated in the Experiment 1 (2 males, one the author and 6 females; age range 24-35), and four subjects in the Experiment 2 (2 males and 2 females; age range 32-38); all subjects were compensated 10 € per hour. All had normal or corrected-to-normal vision and gave their informed consent to perform the experiments. The study was conducted in accordance with French regulations and the requirements of the Helsinki convention. All participants had prior experience with psychophysical experiments and (except the author) were naïve to the specific purpose of the experiment.

### Setup

Participants sat in a quiet, dark room. We recorded the right-eye gaze position with an SR-Research Eyelink 1000 desktop mount, at a sampling rate of 1 kHz. In the Experiment 1 participants had their heads positioned on a chin rest, with adjustable forehead rest, at 180cm in front of a white projection screen. A PROPixx DLP LED projector placed behind the participants and above their head was used to project the stimuli onto the screen at 120Hz. The image projected onto the screen was 137.5cm wide (resolution 1280×720), covering *≈*42degrees of visual angle (dva). The same setup was used in the Experiment 2, except that the chin rest was now placed at 120cm from the projection screen (which then covered *≈*60dva), and the resolution of the image was 1600×900 (the stimuli were scaled to cover the same area in retinal space). An Apple computer running MATLAB (Mathworks) with the Psychophysics and Eyelink toolboxes controlled stimulus presentation and response collection in both experiments.

### Stimuli

Stimuli were noise patterns presented within a Gaussian contrast envelope moving at 12 dva/sec. The noise pattern within the envelope could either drift in a direction orthogonal to that of the envelope (double-drift stimulus), or vary randomly with no net motion (control stimulus). Double-drift stimuli were generated by taking square snapshots from a 2D surface of 1*/f* luminance noise. In the Experiment 1 each snapshot was shifted 2 pixels from the previous one along a constant direction, resulting in a speed of *≈*8 dva/sec. Control stimuli were generated by taking square 2D snapshots from a 3D volume of 1*/f* noise, with each snapshot shifted in depth by 1 pixel from the previous one (this value was chosen to match the perceived temporal frequency of the double-drift stimulus on the basis of preliminary tests; the results of the control task confirmed a posteriori that this value made the two stimuli perceptually similar when seen in the periphery). Since the 3D noise volume was anisotropic, the internal pattern in the control stimulus varied but did not contain any prevalent direction, and therefore differently from the double-drift stimulus it did not induce any systematic distortion of the motion direction. In the Experiment 2, double-drift stimuli with different internal speeds were obtained by shifting the noise pattern by 1, 2 or 3 pixels (corresponding to *≈*5, *≈*10 and *≈*15 dva/sec, respectively); control stimuli had also 3 different temporal rates, created by taking subsequent 2D snapshots in depth at half the rate of the double-drift stimulus (obtained by interpolating the noise pattern for distances of half pixel). Each snapshot was multiplied with a 2D Gaussian with SD set to 0.35 dva, and constituted 1 frame (*≈*8.3 ms) of stimulus presentation. The noise patterns were generated randomly at the beginning of each trial.

### Procedure

#### Experiment 1: main task

Each trial began when the participant’s gaze position was detected within a circular area of 2 dva of diameter centered on the fixation point (a black disk of 0.2dva of diameter) continuously for more than 200 ms consecutively. After a random interval (uniformly distributed within 600-1000 ms) the stimulus appeared at a random position, moving either toward or away from fixation at 12 dva/sec. The starting position of the target was defined on each trial by drawing a random angle (uniformly distributed in (0, 2*π*]) and adjusting the radius so that the target would (given its direction, inward vs. outward) reach an eccentricity of 8dva at the time of saccade landing. The expected time required by the participant to complete the saccade was estimated on every trial by summing the average saccade latency of the previous 20 trials (assuming a value of 200 ms for the first 20 trials) with the expected duration of the saccade, which we estimated as 36 ms. The duration was estimated according to the formula 23.6 + 2.94*A*,Abrams et al. (1989), *where A* is the saccadic amplitude and assuming a 10% amplitude undershoot Becker (1989); Lisi et al. (2019). The moving target appeared and immediately started moving and the participant was required to intercept it with a saccadic eye movement (see Fig. 1).

**Figure 1:**
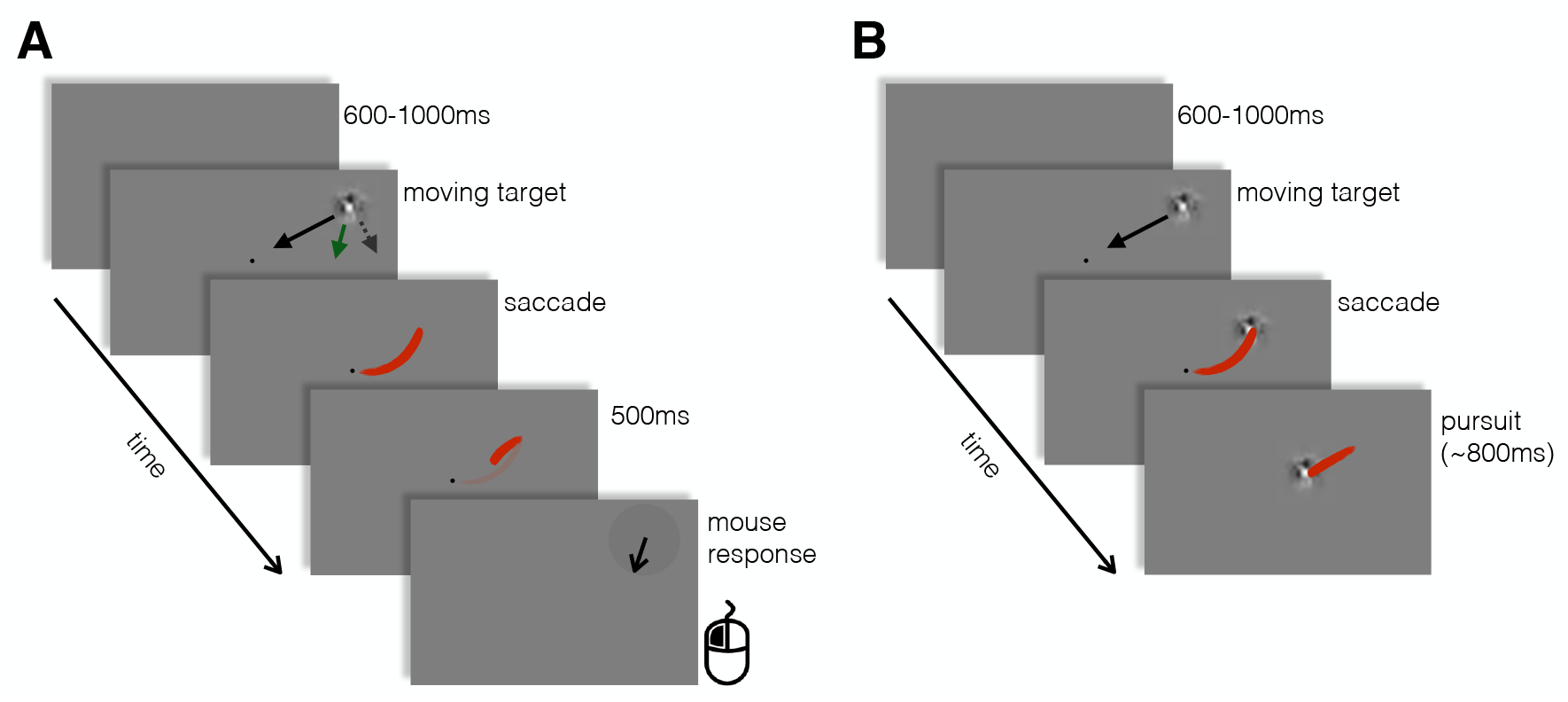
Experimental protocol. A. Trial sequence in the double-drift condition. The red lines represent hypothetical gaze position traces, during and after the saccade. Even when the target was not displayed after the saccade landing, pursuit was initiated after the landing Krauzlis and Lisberger (1994). The black line indicates the physical motion of the envelope on the display, the dashed arrows indicate the direction of the internal motion, and the green arrow the hypothetical perceived direction. At the end of the trial the participant reported the direction of the moving target. B. Trial sequence in the control condition. Here the target remained visible after the saccade and participants had to track it until it disappeared. These two conditions were randomly interleaved, and difficult to distinguish prior to the saccadic response (verified by a subsequent control condition). Participants were never asked to report the perceived direction of the target in control condition; the purpose of the control condition was to provide a baseline for eye movement analyses and, by providing post-saccadic vision of the moving target, to make pursuit a default response upon saccade landing.

Unbeknown to the participants on half of the trials the target was a double-drift stimulus (double-drift condition) whereas in the remaining half, randomly interleaved, the target was a control stimulus (control condition). In addition the different target stimulus (see Stimuli for details), the trial sequences of these two condition diverged after the saccade landing. On double-drift trials the target was removed during the saccade, and was replaced after 500ms by a black arrow (random orientation); in these trials the participants were asked to report the direction of motion of the moving target by adjusting the direction of the arrow with the mouse. Importantly, even though there was not target present after saccadic offset in these trials, and therefore no visual motion signals, we expected to find a short (*≈* 100ms) post-saccadic pursuit response, corresponding to the open-loop response of the pursuit system Krauzlis and Lisberger (1994), which must have been driven by pre-saccadic velocity information.

On control trials instead participants were not required to report the perceived direction. The target was a control stimulus (i.e. containing no internal motion, see Stimuli for details) that remained visible and continued moving after the saccade; participants were required to track the target with their gaze until it disappeared. The purpose of the control condition was mainly to provide a baseline for eye movement analyses (in particular for saccade analyses), and to avoid that repeated post-saccadic disappearances of the target resulted in an attenuation of the post-saccadic, open-loop pursuit response. Indeed, control and double-drift stimuli were designed to appear as similar as possible, such that participants would not realize that two qualitatively different types of stimuli were being presented. To test that this manipulation was successful, after the main task we revealed that there were two types of stimuli and measured participants’ ability to discriminate the two types of stimuli (see Experiment 1: control task for details).

The instrutions given to participants were to saccade and track the moving object with their gaze and that, if the object disappeared, they would have been required to report its direction of motion by adjusting an arrow. As mentioned in the Stimuli section, we designed control and double-drift stimuli so that the perceived temporal rate of change of their internal textures was as similar as possible. Furthermore, we introduced random deviation in the trajectories of control stimuli to make them similar to the trajectories of double-drift stimuli. Indeed, although the trajectories of double-drift targets were always physically aligned on a radial direction with respect to fixation, the perceived trajectories were shifted toward the direction of the internal motion Kwon et al. (2015); Lisi and Cavanagh (2015); Shapiro et al. (2010); Tse and Hsieh (2006). In order to make the trajectories of control stimuli as similar as possible to the trajectories of double-drift stimuli, we introduced random shifts in their direction angle: specifically, we rotated them around the expected interception point (at 8 dva eccentricity) by either adding or subtracting (with equal probability) an random angle uniformly distributed within 15-35°(this range was defined to includes perceptual shifts observed in pilot tests with the same paradigm).

Trials in which participants blinked or moved their gaze from fixation before the appearance of the target were aborted and repeated within the same block. Each participant performed 2 sessions of the task, each comprising 400 trials split in 4 blocks. Participants were allowed to take a break whenever they felt the need. A standard 9 points calibration procedure for the eye movements was repeated after each break.

#### Experiment 1: control task

Given the short pre-saccadic stimulus presentation, it was difficult to distinguish between the two types of stimuli before the saccade. Indeed, after being debriefed, all the naïve participants reported not having noticed any difference in the stimulus between the two trial types (that is between trials in which they should track the target after the saccade and trials where the stimulus disappeared after the saccade and they made a direction response). To make sure that the two condition were not distinguishable, we asked each participant to run a subsequent control experiment. We explained to the participants that in the previous task there were two different motion stimuli (control and double-drift) by showing static examples of each. Next we presented stimuli (generated with exactly the same procedure and parameters as in the main task), and asked them to maintain fixation and report, after the stimulus disappeared, whether the stimulus was a control or a double-drift stimulus (no saccade or pursuit required). The stimulus was presented for a variable duration, and after it disappeared two static examples of the stimuli were presented on the left and right side of fixation; participants provided the response by pressing either the left or the right arrow. The duration of the stimulus was adjusted online according to a weighted up-down staircase procedure that sought to find the duration of the target that yielded 75% correct responses. There were 4 independent staircase procedures, 2 for the inward direction and another 2 for the outward direction, starting at 100 ms and 800 ms of duration. The staircases were randomly interleaved, and each of them comprised 80 trials.

### Experiment 2

Experiment 2 used the same procedure as the main task of Experiment 1, with the only difference that 3, randomly interleaved, internal speeds or temporal frequencies were used. Each participant performed 4 sessions of the task, each comprising 480 trials split in 8 blocks.

### Analysis

In the main task of Experiment 1 and in Experiment 2, we analyzed perceptual response and eye movement data to give, for each trial, three independent measures of the effect of the internal motion (in the double-drift condition): one from saccade landing positions, the second from the direction of the post-saccadic pursuit, and the third one from the perceptual reports (see Fig. 2). For each of these measures, we computed the angular differences between the reported/estimated direction, and the direction of the displacement of the target. To obtain a measure of direction bias induced by the internal motion, we transformed these differences so that positive value would indicate a deviation from the physical direction (external motion) in the direction of the internal motion (see Fig. 3B). Examples of the distributions of angles obtained are reported in Fig.. 4A for two of the participants.

**Figure 2:**
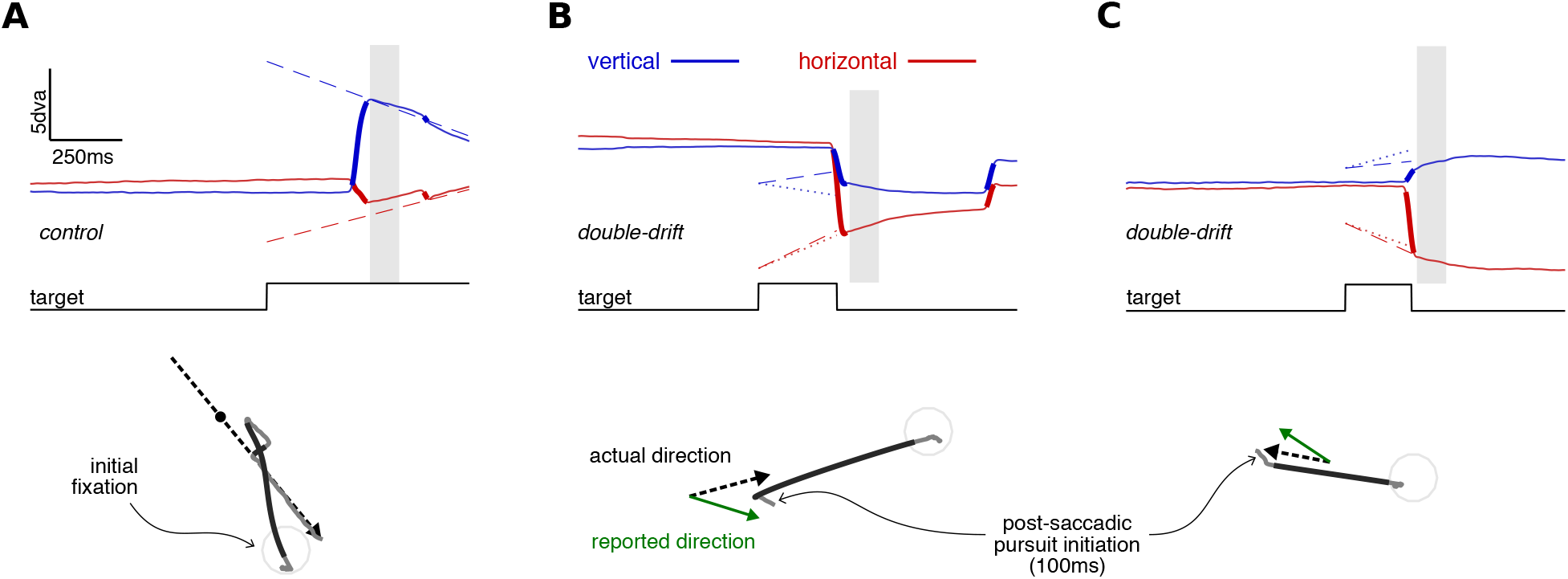
Example of eye movements traces (Experiment 1). A. Example of a control trial: on the top panel the horizontal (red) and vertical (blue) gaze (continuous line) and target (dashed lines) positions are plotted as a function of time. The thicker parts of the gaze position traces indicate saccades. On the bottom panel, the same trial is represented in x-y screen coordinates: the dashed black line indicates the trajectory of the target (the small dot indicates the position of the target at saccade landing time) and grey lines indicate gaze position (the darker parts indicate saccades). In this trial the target moved inward from a peripheral position; the participant first intercepted it with a saccade and then tracked until it disappeared. The vertical grey square delimits the temporal window used for pursuit analysis (20-100 ms after saccade landing time). B-C. Examples of double-drift trials. Notice that the target disappeared after saccade onset, but nevertheless pursuit was initiated after the saccade landing. The dotted lines in the top panels, and the green arrow in the bottom panels represent the direction that the participant reported at the end of the trial.

**Figure 3:**
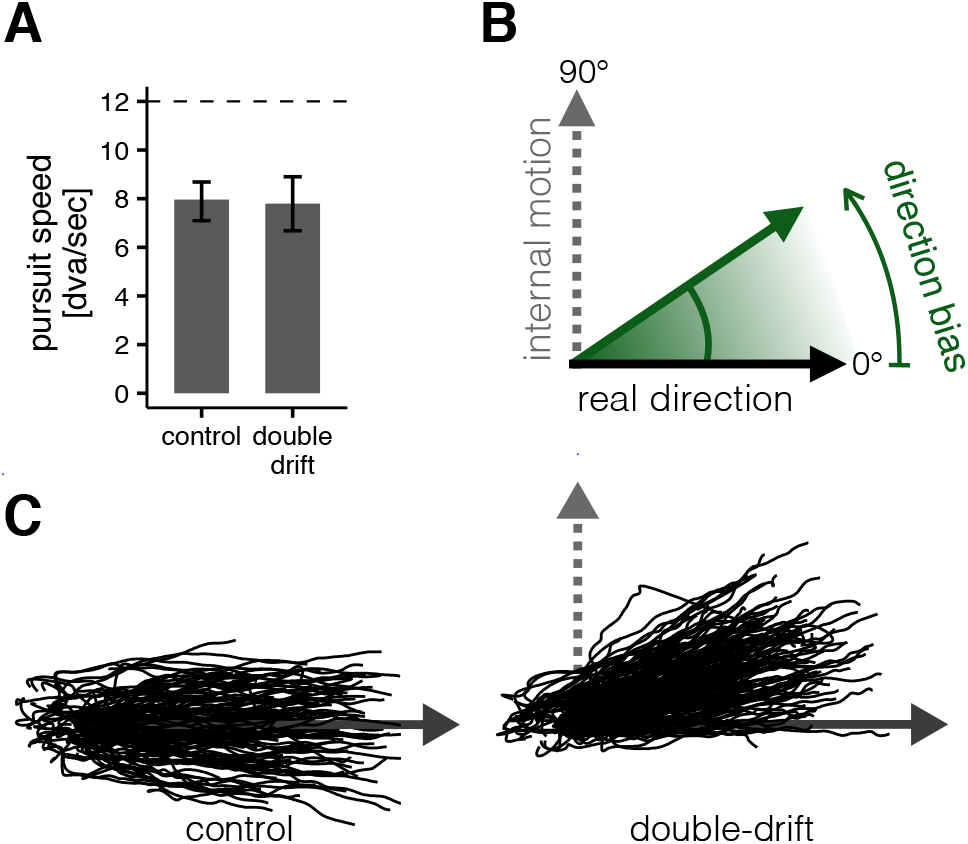
Experiment 1, data analysis: post-saccadic pursuit traces. A. The average speed of pursuit in the post-saccadic interval (20-100ms after saccade offset) was similar independently of the condition (control vs. double-drift) and thus independently of whether a moving target was actually displayed or not. Error bars indicate 95% CI. B. Computation of the direction bias. Each post-saccadic pursuit trace was rotated so that the real direction of motion of the target was at 0° and the direction of the internal motion was at 90°; the same convention was used for the analysis of the direction bias in perception and saccade landings. C. Raw post-saccadic gaze position traces from one participant (rotated according to the conventions illustrated in panel B); in the double-drift condition, the whole distribution of traces is pulled from the real direction of motion toward the direction of the internal drifting pattern.

**Figure 4:**
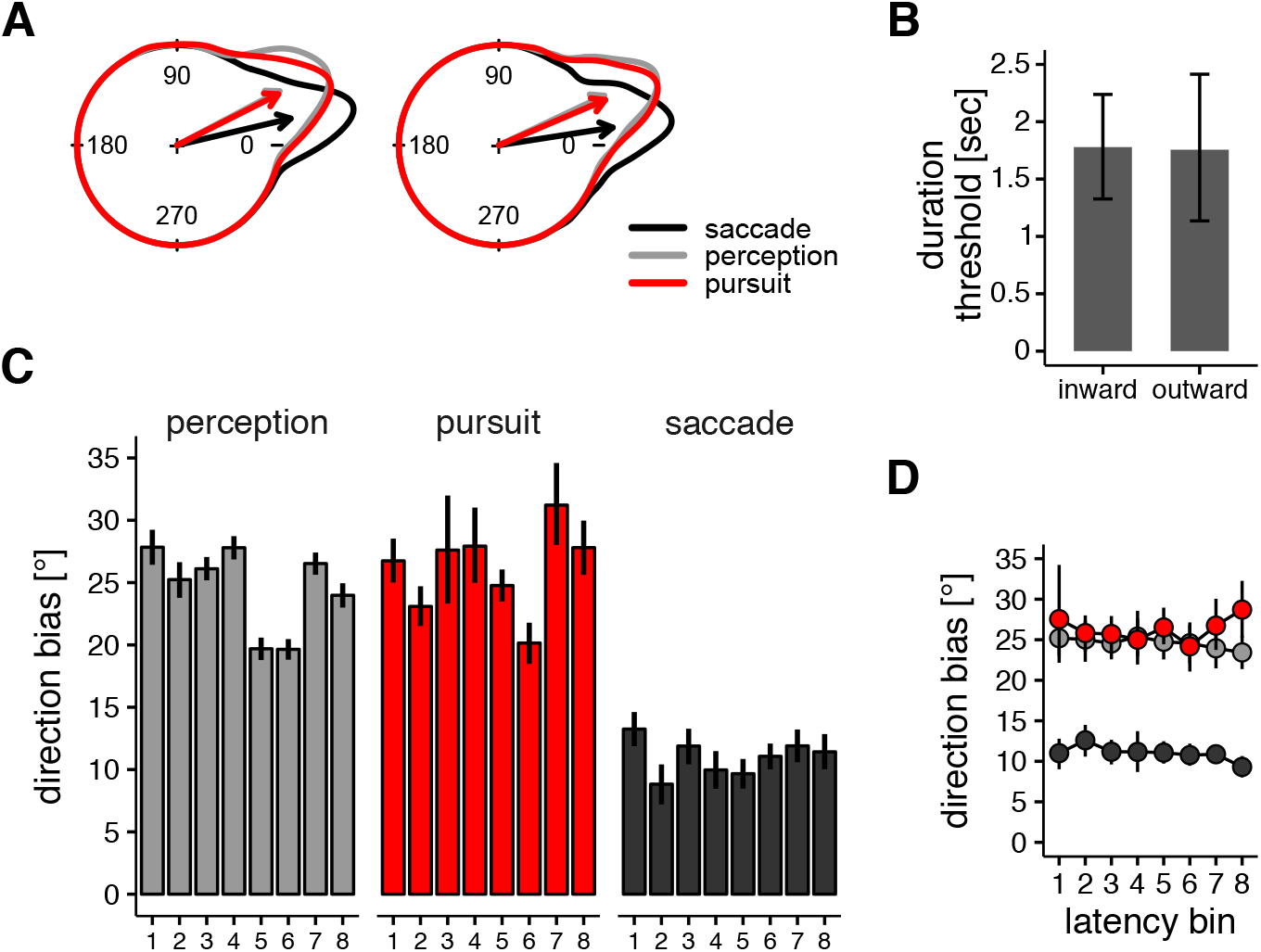
Experiment 1: Results. A. Examples of individual distributions of direction bias (same conventions as figure 3B) for participants 1 and 2. Arrows indicate mean direction biases. B. Results of the control experiment. The duration threshold is the minimum duration of presentation required to reliably discriminate (75% correct discriminations) the double-drift stimuli from the control stimuli in the fixation condition. Notice that the thresholds are much longer than the presentation duration in the main experiment (max 600ms) indicating that participants would be at less than threshold discrimination levels in the main experiment. C. Direction biases for all 8 participants. Error bars represents bootstrapped 95% CI (percentile method). D. The size of the directional bias was not modulated by the latency of the movement for either saccade, pursuit, or perceptual responses (see text for details).

### Deriving the saccade system’s estimate of stimulus speed and direction

Raw gaze position data were transformed to velocities and smoothed by using a moving average over 5 data samples Engbert and Kliegl (2003); saccade onsets and offsets were detected using an algorithm based on 2D eye velocity Engbert and Mergenthaler (2006). Next, we estimated the direction of motion that was used by the saccadic system to intercept the target in double-drift condition. We computed a metric known as saccadic velocity compensation, SVC Groh et al. (1997) by computing the difference between the spatial position of the saccade end point and the starting position of the target, and dividing it by the temporal interval from target onset until the end of the saccade. The SVC corresponds to the target velocity vector (speed and direction) for which the saccade would have been maximally accurate (i.e. for which the landing error would have been zero). The variability of this measure is mostly contributed by the typical scatter of saccade landing positions van Opstal and van Gisbergen (1989), but might also reflect eye tracker inaccuracies, direction estimation biases Krukowski and Stone (2005) or other systematic motor biases. To correct systematic biases and improve the accuracy of the estimated SVC, we calibrated our analysis using control trials, where there was no internal pattern motion to distort the direction that would be taken into account by the saccadic system. For each participant and direction condition (outward vs. inward), we fit a number of multivariate linear models, with the sine and cosine of the target direction angle as dependent variables. In all, we tested 32 different models; all of them included as a predictor in a trigonometric polynomial for the SVC direction angle (of degree 1 or 2, depending on the model), plus one or more of the following variables: direction of the saccade (also represented through a trigonometric polynomial of degree 1 or 2), with or without interaction with the SVC, and interactions between the SVC components and the saccadic latency or amplitude. These models correspond to a circular-circular regression Jammalamadaka and SenGupta (2001) with multiple predictors. To avoid over-fitting, we evaluated the out-of-sample predictive performance of each of the 32 models through a leave-one-out cross-validation procedure (see Figure S1). The best model selected according to this procedure is the one that is expected to generalize better to an independent dataset Geisser (1975), so we used it to estimate the target direction taken into account by the saccade system in the double-drift condition. The final model included as predictor the raw direction angle of the SVC vector in interaction with a trigonometric polynomial of degree 2 of the saccade direction angle. This latter component improved the fit of the model by accounting for saccade biases, which are known to vary as a function of saccade direction Rolfs et al. (2010). The average cross-validated error of the best model (measured in control trials) was 11.36°, about half of the average direction error of the raw SVC measure on the same trials, 20.03°.

### Estimating post-saccadic pursuit direction and speed

Post-saccadic gaze position traces were low-pass filtered using a 100-points finite impulse response (FIR) filter with cut-off at 40Hz. We restricted the analysis to pursuit initiation, from 20 ms to 100 ms after the landing of the first interceptive saccade (we excluded the first 20 ms after saccade landing to prevent any contamination of our pursuit measures by potential inaccuracies in the identification of saccade offsets). This is within the time range of typical open loop pursuit Krauzlis and Lisberger (1994), implying that in this time interval post-saccadic visual signals have not yet had time to influence the ongoing pursuit. We fit these post-saccadic position traces with a line (orthogonal regression) to estimate the direction of the pursuit. Additionally, we differentiated these position traces to obtain a velocity trace, which we averaged over the whole 80 ms interval to get an estimate of the horizontal and vertical pursuit speeds. We took the Euclidean norm of this velocity vector as a measure of pursuit speed (irrespective of direction), which we used to compare the pursuit gain in the two conditions, control and double-drift (see Results). We used the same procedure to measure gaze velocity before the saccade, within an interval from 80 ms to 20 ms before saccade onset, to investigate whether we could detect pre-saccadic pursuit initiation Lindner and Ilg (2000) in our paradigm.

### Perceptual estimate of stimulus direction

We took the direction of the arrow, as set by the participants in double-drift trials, as the perceptual estimates of target direction. For each participant and condition, the size of the perceptual bias was computed by taking the angular mean of the signed angular differences between the arrow and the true direction of motion of the target.

### Control task

We analyzed the proportion of correct identifications of the type of stimulus (double-drift vs. control) as a function of the duration of presentation, to determine the minimum duration required to achieve 75% of correct identifications. We fit the data separately for each direction (inward vs. outward), using a generalized linear mixed-effects model with a probit link function and a fully parametrized variance-covariance random effect matrix, fitted with R R Core Team (2015) and the lme4 Bates et al. (2012) library.

## Results

### Experiment 1

#### Perception and pursuit reveal similar estimates of target velocity

The offline analysis of eye movements excluded 4% of the trials, due either to missing values in the gaze position trace, or to the execution of a blink or an eye movement before the appearance of the target. Of the remaining trials, 0.51% were excluded because the saccade latency was either shorter than 100ms or longer than 600ms. Additionally, 0.78% of the trials were excluded due to extreme saccade amplitude values, either less than 3dva or larger than 20 dva.

In the remaining trials we found that, on average, the perceived direction reported by the participants was shifted by 24.54° (SD 3.69) from the physical direction toward the direction of the internal motion. This shift in the mean direction was significantly different from 0 for each participant (see Fig. 3), but it was about 25% smaller than what would be expected according to the simple sum of the two motion vectors (*≈*32°). There was no significant difference between the bias measured in trials with inward vs. outward motion in the target, *t*(7)=1.65, *p*=0.14.

To examine how the pursuit system responded to the double-drift stimulus, we analyzed the early post saccadic pursuit initiation. The speed of the pursuit from 20 to 80 ms after saccade landing was on average 7.88 dva/sec (SD 1.44), corresponding to a gain of 0.66 from the stimulus speed of 12.0 dva/sec. The average speed in the control condition was 7.96 dva/sec (SD 1.23) and 7.79 dva/sec (SD 1.69) in the double-drift condition. A repeated measures ANOVA revealed that pursuit speed was not influenced by the condition (double-drift vs. control), *F* (1, 7)=0.66, *p*=0.44; nor by the direction of motion (inward vs. outward), *F* (1, 7)=0.65, *p*=0.45; nor by the interaction between condition and direction, *F* (1, 7)=2.41, *p*=0.16. This finding reveals that the eye movement behavior did not differ, in the first 100ms after saccade landings, between the control and double-drift conditions, despite the fact that the moving target was present only in the control condition (in the double-drift condition the screen was blank after the saccade). We then restricted the analysis to double-drift trials. We found that the direction of the pursuit was shifted on average by 25.76°(SD 3.74) from the physical direction of motion of the target, toward the direction of the internal motion. The size of the shift did not differ across inward vs. outward trials, *t*(7)=1.64, *p*=0.14. The deviations of direction measured in pursuit were not different from those measured in the perceptual responses, *t*(7)=1.13, *p*=0.29 (see Fig. 4C). Additionally, there was a tendency for a positive correlation across participants between pursuit and perceptual responses. The correlation was statistically significant with a one-tailed test, *r*(7)=0.63, *p*=0.04. We assessed also whether there were measurable correlations on a trial-by-trial basis. Correlations were quantified using a measure of circular dependence which has the same properties of the product moment correlation coefficient for linear variables, but is appropriate for angular variables Jammalamadaka and Sen-Gupta (1988, 2001). The correlation coefficients resulted in: perception-pursuit, mean 0.10, range from 0.02 to 0.18; saccade-pursuit, mean -0.09, range from -0.14 to -0.02; saccade-perception, mean 0.02, range from -0.07 to 0.09. Of these, the only trial-by-trail correlations that were statistically significant (at the Bonferroni-corrected Type 1 error rate *α*=0.00625) were the positive correlations between perception and pursuit responses, for 3 out of 8 participants.

#### Velocities underlying saccadic extrapolation are not consistent with pursuit and perceptual velocity estimates

Here we use the saccade landings to derive the estimate of stimulus velocity used by the oculomotor system to program the saccades. The average saccadic latency in the trials included in the analysis was 226 ms (standard deviation across participants, SD, 16) in control trials, and 230 ms (SD 20) in double-drift trials; the latency did not differ across these two conditions, *t*(7)=1.38, *p*=0.21 (paired t-test).

In order to best estimate the target direction used by the saccadic system, we fit a model suited for angular data on the control condition, validated it through a cross-validation procedure (see Supplemental Figure 1), and then used it to estimate for each trial what was the most likely target direction that elicited the observed saccades in the double-drift condition. The models were fit, on average, with 373 control trials per participants and were used to predict the direction of the target in (on average) 374 double-drift trials. With respect to control trials, the average r-squared of the fit was 0.95 for horizontal (cosine) components and 0.92 for vertical (sine) components. In control trials, the estimated direction was on average shifted by 10.98° (SD 1.48) from the physical direction toward the direction of the internal motion, with no significant difference between trials with inward vs. outward motion direction, *t*(7) = 0.85, *p* = 0.42. This level of direction bias was similar, although slightly smaller, than the one obtained by considering raw SVC measures, 13.55° (SD 2.96). Importantly, saccade responses were significantly less influenced by the internal motion than both perceptual, *t*(7) = 12.20, *p <* .0001, and pursuit responses, *t*(7) = 12.11, *p <* .0001 (Fig. 4C).

We computed the absolute distance between the onset position of the target and the landing position of the saccade, divided by the temporal interval between target onset and saccade landing, and took it as a measure of the target speed (as opposed to direction). The average speed of the target estimated from saccade landing was very similar across control and double-drift conditions: 12.10dva/sec (SD 0.89) and 11.78 dva/sec (SD 1.01), respectively, both very close to the actual target speed of 12.0 dva/sec.

Finally, we investigated whether the observed difference between pursuit, perception and saccades could be related to differences in the latency (see Fig. 4D). We split the dataset in 8 equally sized bins according to the individual saccade latency distributions. Thus, for each participant, each latency bin comprised 12.5% of his or her data ordered according to increasing saccade latency. For each response, we performed repeated-measures ANOVA with the observed direction bias as dependent variable and the latency bin as independent variable. This analysis failed to provide any evidence for modulation of the directional bias by latency in perceptual responses, *F* (7, 49) = 1.12, *p* = 0.36; pursuit responses, *F* (7, 49) = 0.96, *p* = 0.47; or saccade responses, *F* (7, 49) = 1.18, *p* = 0.33.

#### Pre-saccadic pursuit

In Rashbass’s original studies Rashbass (1961), an initial pursuit response could be seen just prior to the saccade. To analyse any pre-saccadic pursuit here, we included in the analysis only trials where the speed of gaze movements in the pre-saccadic interval differed by more than 2 standard deviations from the gaze velocity measured in a baseline interval (300 ms before the onset of the target). The pre-saccadic gaze velocity exceeded this threshold only on 12% of the trials. We computed the difference between the direction of the pre-saccadic pursuit and the direction of displacement of the moving target, and performed a series of Rayleigh’s tests Mardia (1972) to assess whether these angular differences were concentrated along one direction or were uniformly distributed. The null hypothesis of circular uniformity was rejected (at the Bonferroni adjusted level *α*=.00625) in 4 cases out of 8 in the control condition, and in 5 cases in the double-drift condition. Overall, the average direction of the pre-saccadic pursuit resulted roughly aligned with the external direction of the target envelope, in both the control and double-drift conditions (see Fig. S3), however due to the small number of trials and the large variability it was not possible to determine whether the pre-saccadic pursuit was also significantly biased by the internal motion.

#### Perceptual similarity of control and double-drift stimuli

In the main experiment, the control trials, where the target remained after the saccade, were intermixed with the double-drift trials, where the target was absent after the saccade. The goal of this procedure was to encourage participants to make pursuit responses in the double-drift trials. In order for this strategy to be effective, the control and double-drift trials need to be indistinguishable. To examine this, we ran a control experiment in which we assessed the minimum duration of stimulus presentation required for the participants to reliably discriminate the two types of stimuli (control and double drift; see Stimuli section). Overall, distinguishing the two types of internal motion was quite difficult, and the duration yielding 75% of correct responses was much longer than the average saccadic latency (or duration of presentation) used in our main task (see Fig. 4B). More specifically, the 75% threshold was estimated in 1778 ms, bootstrapped 95% CI [1326, 2238], for inward moving targets, and 1755 ms, bootstrapped 95% CI [1135, 2414].

### Experiment 2

#### Varying internal motion influences only estimated target direction, not speed

In Experiment 2 we varied the speed of the internal drift, to gain some insight into how the two motion signals are combined. To analyze Experiment 2, we used the same rejection criteria as in Experiment 1, which led to the exclusion of 1.9% of trials due to saccade latencies either shorter than 100 ms or longer than 600 ms, and to the exclusion of 0.2% of trials due to saccade amplitudes shorter than 3dva or larger than 20dva. The average saccadic latency in the remaining was 219 ms (SD, 42 ms) in control trials, and 209 ms (SD 35 ms) in double-drift trials. We did not find evidence for a modulation of saccade latency due to the speed/temporal frequency of the noise, *F* (2, 6)=2.85, *p*=0.13, the condition (control vs. double-drift), *F* (1, 3)=4.66, *p*=0.12, nor their interaction, *F* (2, 6)=0.86, *p*=0.47.

The average directions reported by participants were shifted toward the direction of the internal motion by 17° (SD 2°), 29° (SD 6°) and 33° (SD 10°) for the conditions with internal speed set to 5, 10 and 15 dva/sec, respectively (see Fig. 5). The effect of the internal motion speed on the perceptual direction bias was statistically significant, *F* (2, 6)=14.06, *p*=0.005, whereas the effect of target direction (inward vs outward), *F* (1, 3)=7.17, *p*=0.07, and the interaction between direction and internal speed, *F* (2, 6)=3.57, *p*=0.09, did not reach statistical significance. The observed direction biases were again about 25% smaller than what predicted by a simple vector sum of the two motion vectors (*≈* 22°, 36°and 44°). Similarly, the analysis of pursuit responses revealed a strong effect of the speed of the internal motion, *F* (2, 6)=132.5, *p <*0.0001, but no evidence for an effect of direction (inward vs outward), *F* (1, 3)=0.97, *p*=0.40, or for an interaction, *F* (2, 6)=2.01, *p*=0.21.

**Figure 5:**
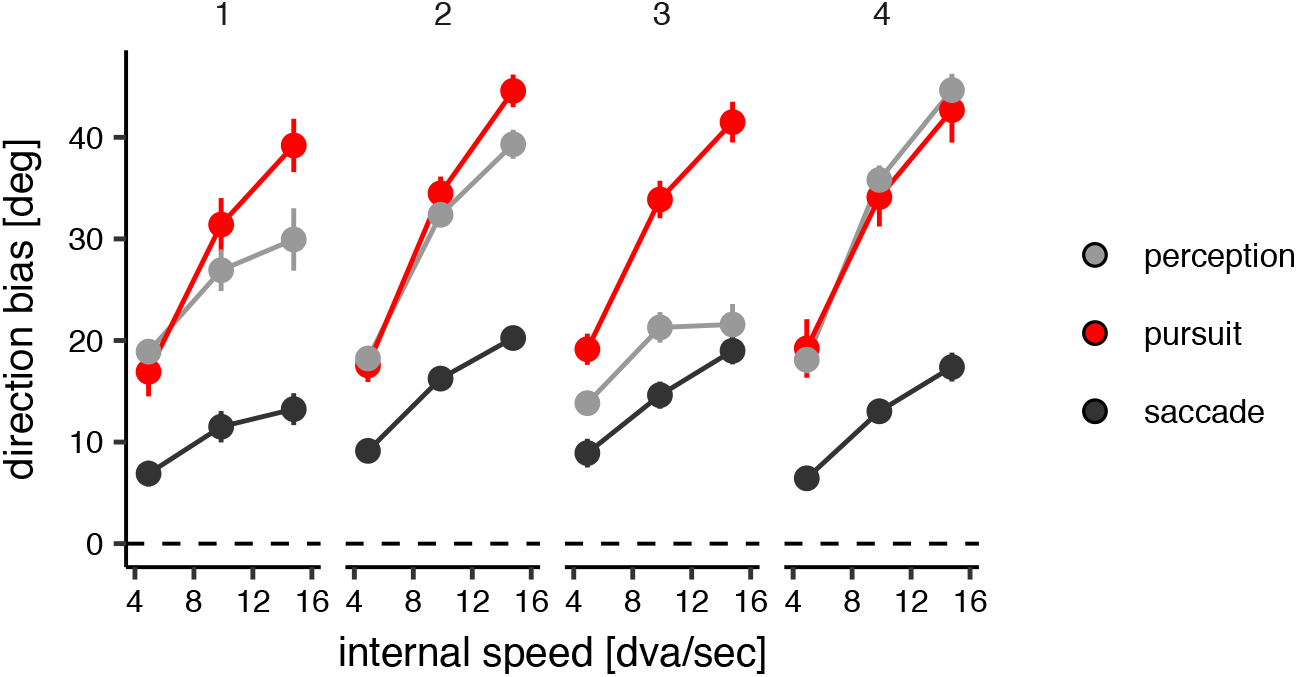
Experiment 2: Results. Direction bias in perception, saccade targeting and open-loop pursuit plotted as a function of the speed of the internal motion. Each panel represents data from one participant. Error bars are 95% confidence intervals.

To analyze saccade responses, we used the same model selected for the analysis of saccade responses in Experiment 1. In this case the model was fit on average on 899 saccades for each participant. The estimated direction biases revealed that saccade landings were also influenced by the speed of the internal motion, *F* (2, 6)=72.87, *p <*0.0001, but not by its direction, *F* (1, 3)=0.30, *p*=0.62, or their interaction, *F* (2, 6)=0.53, *p*=0.62.

We then investigated whether the pursuit speed and the estimate of target speed recovered from saccade landings varied as a function of the speed of the internal motion. Indeed, if the integration mechanism was a vector averaging or vector sum, the increased speed of the internal drift should yield an overall faster speed estimate, and therefore produce higher pursuit speeds and saccade endpoints that are extrapolated even more along the direction of motion. Nevertheless, the three-fold variation of internal drift speed had no significant effect on pursuit speed or the speed estimate of the saccadic system (see Table 1), pursuit, *F* (2, 6) = 2.58, *p* = 0.16, saccades *F* (2, 6) = 0.76, *p* = 0.51.

**Table 1:** Target speed (mean and standard deviations across observers) as inferred from the distance between target onset position and saccade endpoint and speed of post-saccadic pursuit, as a function of the speed of the internal motion.

| Internal speed | Saccades | Pursuit |
| --- | --- | --- |
| 5 dva/sec | 11.90 (0.97) | 6.58 (0.52) |
| 10 dva/sec | 12.18 (1.18) | 6.88 (0.68) |
| 15 dva/sec | 12.06 (1.34) | 6.72 (0.49) |

In sum, the results of Experiment 2 revealed that the internal drift speed had a clear effect on the direction of the responses. However, it did not produce a significant effect on the speed estimates driving the eye movements.

## Discussion

In this study, we investigated the relationship between motion estimates used in saccades, pursuit, and perception. We presented moving apertures filled with noise that either drifted orthogonally to the aperture’s direction (a double-drift stimulus), or varied dynamically with no net motion. The first of these two, the double-drift stimulus, produces a remarkably robust perceptual deviation of direction of up to 45°or more Lisi and Cavanagh (2015); Shapiro et al. (2010); Tse and Hsieh (2006). Using this stimulus we have previously showed a dissociation in the localization of moving objects between saccades and perception Lisi and Cavanagh (2015). This initial report however could not pinpoint whether the dissociation was due to how saccades and perception processed motion signals versus how they used them to extrapolate the position of the moving object. To address this question, in the current study we designed a novel protocol to measure and compare the effects of the illusion in perceptual judgments and two types of eye movements, saccades and pursuit with all three measured on each trial in which a double-drift stimulus was presented. The targets appeared in the periphery and moved either toward or away from fixation. Participants were asked to saccade to these targets as soon as they appeared, and then track them with their gaze. Participants were instructed that in some trials the target would disappear upon saccade landing, in which case they would have been required to report its direction using a directional arrow. In these trials, early post-saccadic pursuit (open loop) still occurred so that the effect of the pre-saccadic illusory direction was seen in the post saccadic pursuit, enabling a trial-by-trial comparison of perception, and pursuit responses. We also estimated the direction and velocity of target motion taken into account by the saccadic system from the saccade landing errors. The results showed that motion signals from the internal drift strongly affected not only the perceived target direction but also, with a similar magnitude, the direction of post-saccadic pursuit. In contrast, the effect on saccade landing positions appeared substantially smaller, less than half the shift observed in perception and in the pursuit traces.

In the Experiment 2, we varied the speed of the internal drift motion and observed changes in the direction of the pursuit, but not in its speed. This indicates that the oculomotor system combines the internal and external motion components of the double-drift stimulus according to some integration process that does not conform to either simple vector averaging or vector summation. Saccadic responses showed a similar pattern. We calculated the target speed estimated by the saccade system by observing the offset of the saccade landing from the initial position of the target (see Analysis section). The saccade system compensates for target motion during the delay of programming and execution of the saccade by extrapolating the landing along the direction of target motion Fleuriet and Goffart (2012); Fleuriet et al. (2011). We found that changing the internal speed of the target only influenced the direction of extrapolation, but not the amount of extrapolation. Previous studies have examined the effect of MT microstimulation on monkey’s oculomotor behavior during the tracking of a moving target Groh et al. (1997) and found evidence for a vector averaging mechanism. In contrast, our results do not support vector averaging, suggesting that injecting a motion signal with MT microstimulation and injecting it with a drift within the moving target produce different effects. So even though the brain might not be able to tell apart motion signals generated by physical displacements and those generated by MT microstimulation, it can keep separate velocity estimates for internal (pattern drift) and external (object displacement), as postulated by a recent Bayesian object tracking model Kwon et al. (2015).

The different pattern of errors showed by perceptual judgments and saccadic eye movements that we found here are consistent with the dissociation observed in our previous studies Lisi and Cavanagh (2015, 2017b); Massendari et al. (2018).

Overall, although we find here a small effect of internal motion on saccade landings, the results presented here are consistent with the dissociation

The different pattern of errors showed by perceptual judgments and saccadic eye movements here and in our previous studies Lisi and Cavanagh (2015) could be interpreted as due to a different processing of visual motion signals. For example, sensory maps representing local motion measurements (i.e. monkey areas MT or V5) may be read out for separate tasks according to different algorithms Groh et al. (1997). Indeed, some lines of evidence suggests that visual motion analysis can be specifically tailored for each function it serves Simoncini et al. (2012), so that when confronted with the same motion stimulus, different responses, such as perception and eye movements, can reveal large differences in the estimated parameters (speed or direction) of the motion (e.g. Glasser and Tadin (2014); Spering and Gegenfurtner (2007); Spering et al. (2011). However, we found here that another eye movement response, smooth pursuit, reveals estimates of motion direction that are virtually equivalent to those found for perceptual judgments (see also Maechler et al. (2021)), suggesting that pursuit and perception use similar computation to estimate the velocity vector of the double-drift stimulus. One possible account of the current results therefore could be that saccade and pursuit are informed by different motion processing mechanisms. This however seems unlikely: previous research has consistently shown a tight relationship and cooperative interactions between saccades and pursuit during the tracking of moving targets Erkelens (2006); Fleuriet and Goffart (2012); Goettker et al. (2018); Lisi and Cavanagh (2017a); Orban de Xivry and Lefèvre (2007).

If saccadic eye movements and pursuit share common motion processing mechanisms, how can we explain that the saccade landings were systematically closer to the physical target locations that what would be expected based on the direction of pursuit? One possibility is that the saccadic system uses a similar velocity vector as perception and pursuit, but extrapolates target position for an interval shorter than the saccadic latency. Specifically, the direction bias measured from saccadic landing positions in our task could be consistent with an extrapolation of the target position along the direction of early pursuit for an interval of only about 110 ms (see Fig. S4). This temporal interval is smaller than the typical saccadic latency and approximately compatible with the so-called saccadic dead time, which is the short interval just before the onset of a saccade during which the system cannot make any more changes to the motor command Etchells et al. (2010); Lisi et al. (2022); Ludwig et al. (2007). According to this account, the rapid programming of saccadic eye movements would limit the maximum temporal interval over which the position of the moving target needs to be extrapolated, yielding a smaller position error in the case of the double-drift stimulus. We propose that this could be the result of a dedicated oculomotor map that accumulates information and predicts ahead in time over a shorter interval than perception. In other words, the dissociation between saccadic responses and perceptual judgments would be at the stage where motion signals are integrated with position signals to extrapolate future positions and could be in the length of the temporal interval compensated by the extrapolation. This hypothesis is consistent with the proposal that neural processing across the brain is organized in a hierarchy of temporal scales Heeger (2017); Murray et al. (2014) and could provide a mechanistic explanation to other action-perception dissociations, such as those where eye movements was influenced by perceptually suppressed stimuli Rothkirch et al. (2012). Taken together, these findings support the idea that neural circuits governing where we look process information on a shorter timescale than neural circuits making inferences about what we see Lisi and Cavanagh (2015, 2017b); Massendari et al. (2018).

## Acknowledgments

The research leading to these results has received funding from the European Research Council under the European Union’s Seventh Framework Program (FP7/2007-2013)/ERC grant agreement n°AG324070 to P. C. and from the Dartmouth College Department of Psychological and Brain Sciences to P.C.

---

**Figure S1:**
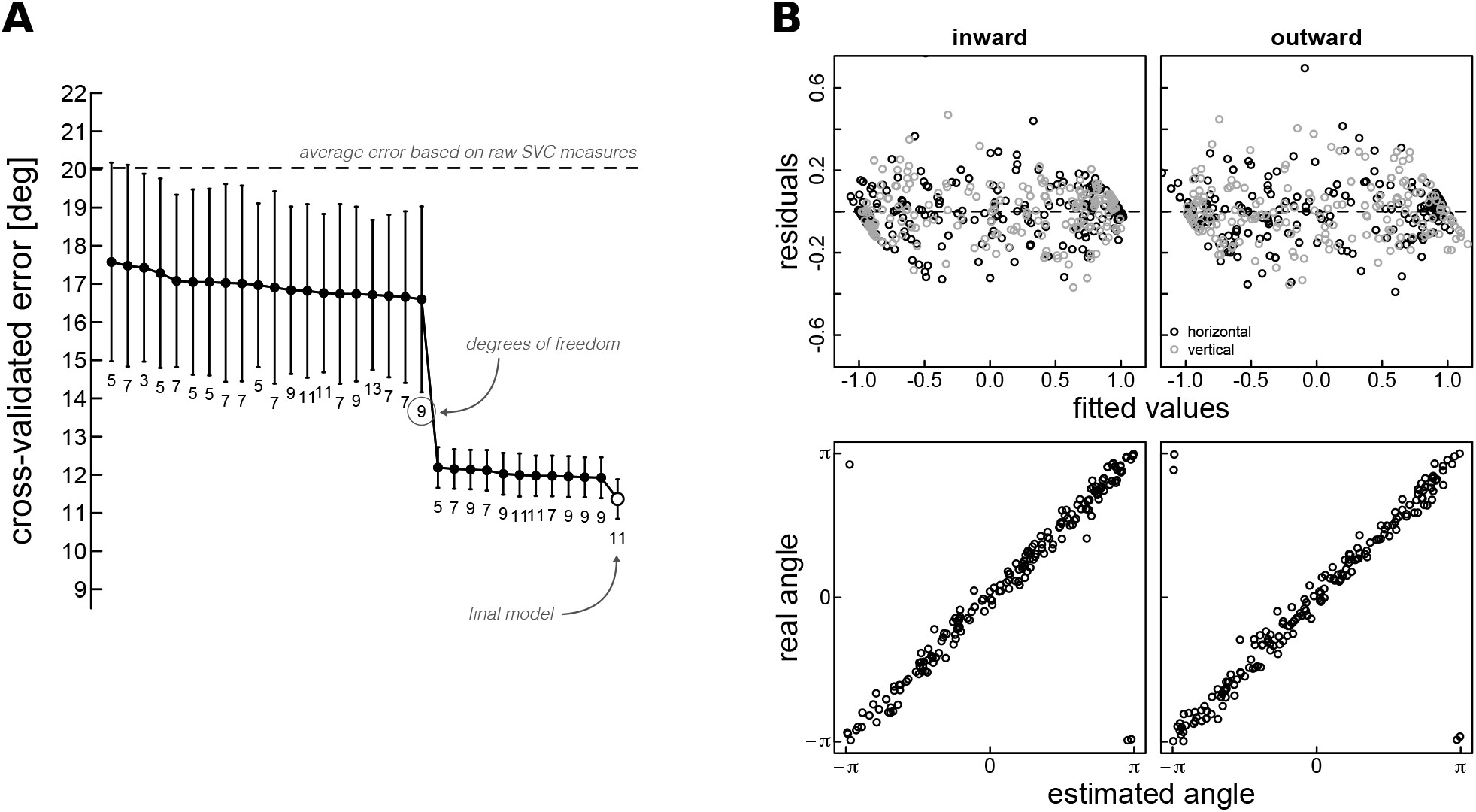
A. Each point represents a different multivariate model; the Y-axis represents the average error on the left-out trials (error bars represents the standard error across participants and conditions) measured in the cross-validation procedure. (Note that the model was fit to control trials where there was no bias on saccadic targeting and therefore we can assume that the direction taken into account by the saccadic system corresponded, except for internal noise, to the physical direction of motion.) This multivariate model was fit separately for each participant and direction (inward vs. outward). The horizontal dashed line represents the average error obtained by using the raw saccadic velocity compensation metrics (SVC) to estimate the target direction. The numbers underneath the dots indicates the number of degrees of freedom for each dimension (vertical and horizontal) of the models. The model used in the analysis presented in the main text is the one that obtained the smallest error and can be formally notated as

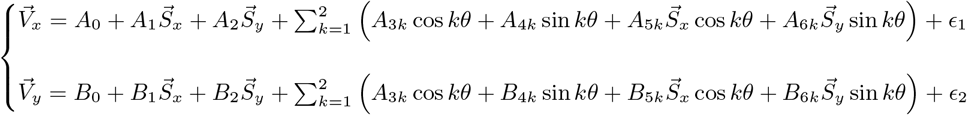

Where 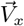 and 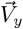are the horizontal and vertical components of the target motion up to saccade landing; 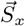 and 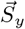 are the horizontal and vertical components of the raw SVC metrics; θ the direction angle of the saccade, and ϵ_1_, ϵ_2_ normally distributed residual errors. We used this analysis to reduce the variance of the estimated trial-by-trial direction errors and increase the sensitivity of our analysis to trial-by-trial correlations between direction errors measured from saccades, perception and pursuit. We point out that analyses performed on raw SVC measures, or with different models all converged in showing the same pattern of results presented in the main text. B. Example of the model fit for one participant. The two panels show the residual error as a function of the predicted velocity components 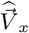 (black dots) and 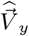 (gray dots); the lower panels show real and estimated values of target direction angle, calculated as arctan2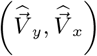(where ‘arctan2’ denotes the four-quadrant inverse-tangent function).

**Figure S2:**
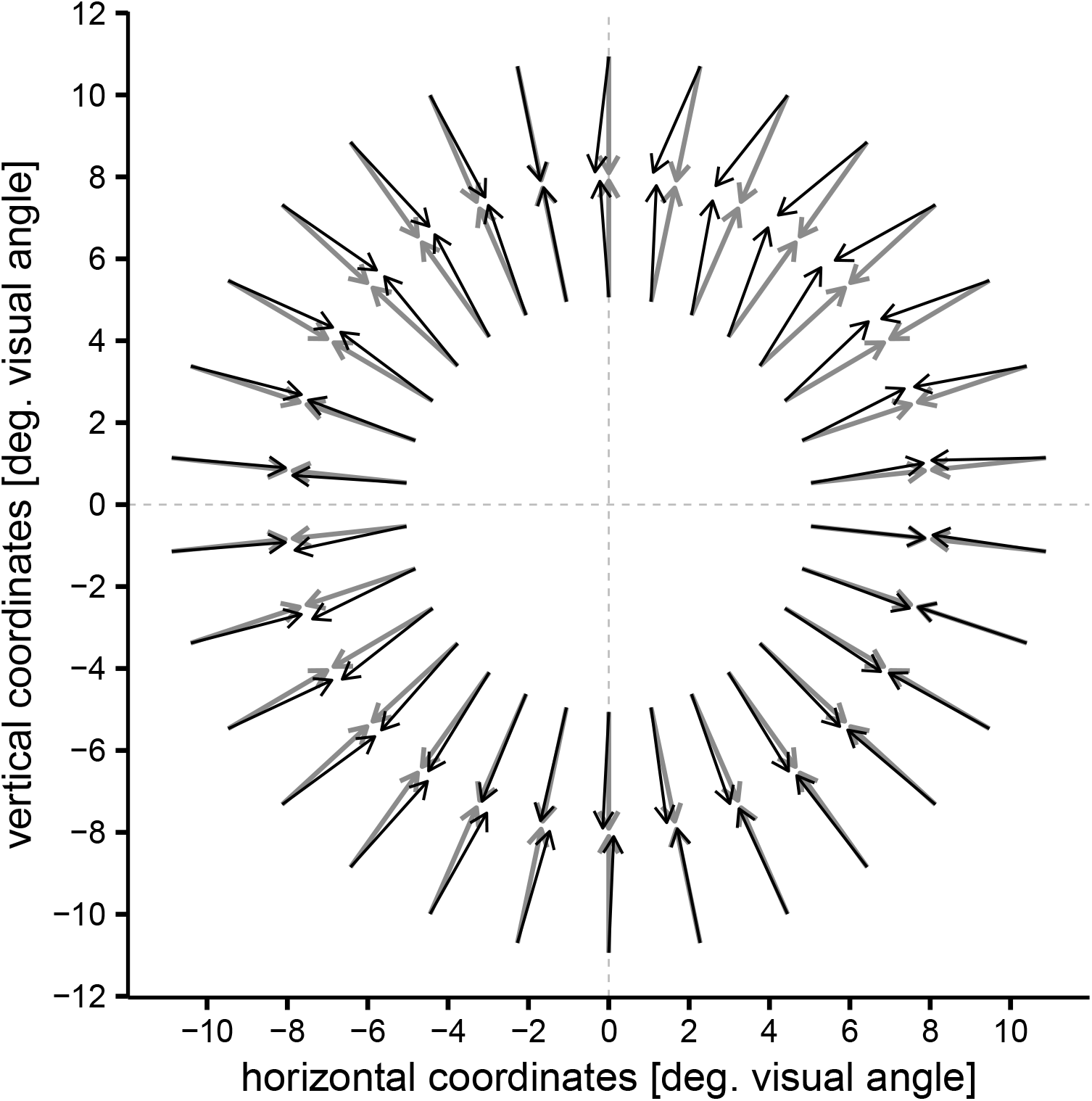
Example of correction of the estimated target direction from saccade landing. Raw saccade velocity compensation (SVC) measures can be affected by idiosyncratic, location-specific biases, possily due to changes in head position after the calibration or imperfect eyetracker calibration. This figure demonstrates how the model used in the analysis of saccade landings (see Main text and Fig. S1 for details) influence the estimated target direction, for one example participant. The grey arrows represents (hypothetical) raw SVC measures, shown for both inward and outward direction, and equally spaced around the visual field. The black arrows represents the predictions of the model, which was cross-validated on control trials to maximize the accuracy in predicting the target velocity vector. Note that the models for inward and outward saccades were fit independently (using data from different trials), yet they are highly consistent in how the vectors are displaced - see for example how the estimated vectors (black arrows) in the top-right quadrant are both shifted toward the left relatively to the raw measures (gray arrows). This suggests that the model-based estimates are correcting a calibration inaccuracy that affected the top-right part of the display, regardless of the target direction (inward vs. outward).

**Figure S3:**
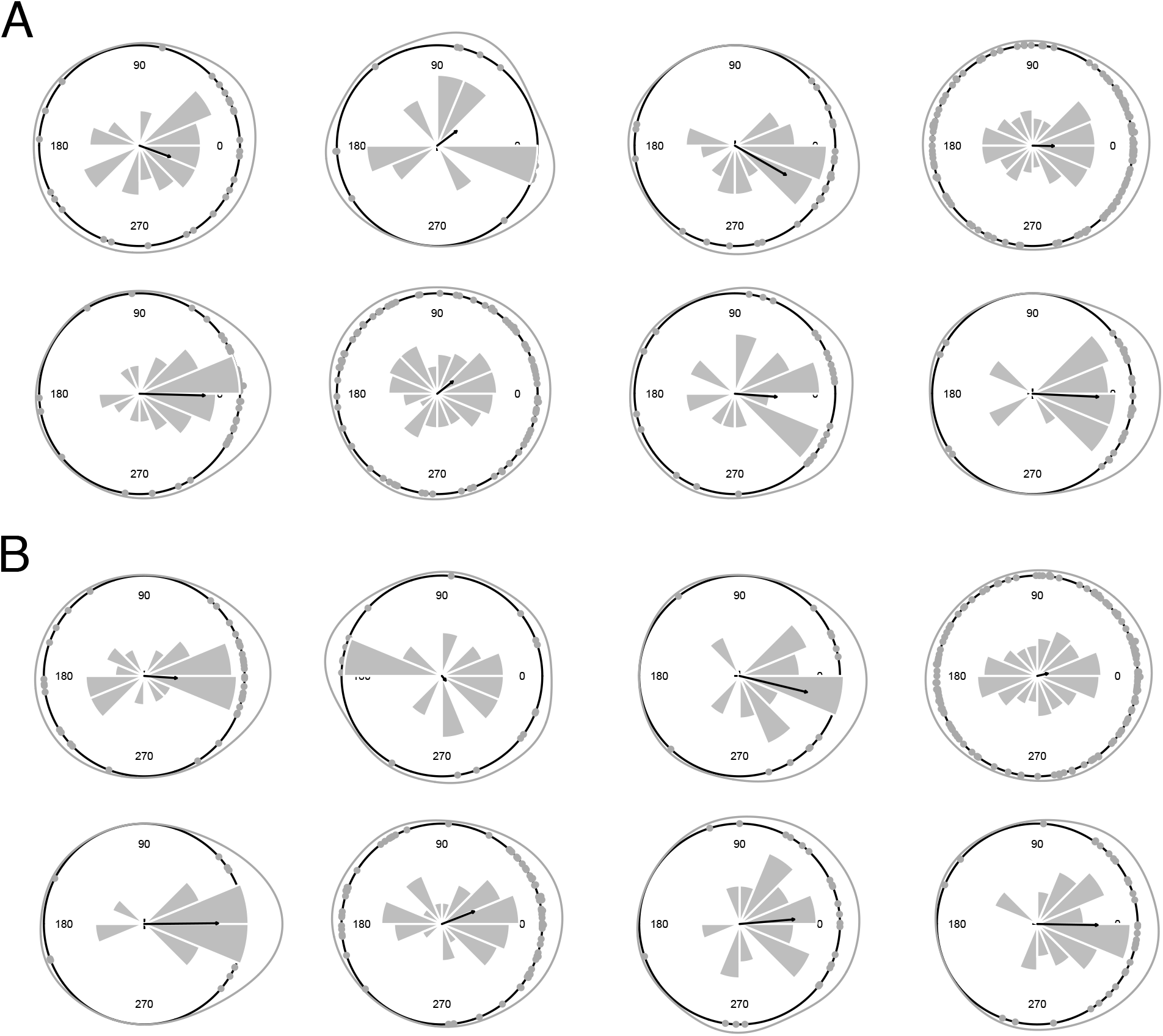
Direction of pre-saccadic pursuit. A. Control condition. Each point in the plots represents the direction of the pre-saccadic gaze movements on a given trial, with respect to the physical direction of the target envelope (which correspond in the plot to 0 degrees); the histogram and the circular density represent the distribution of these points, and the black arrow, the average direction. B. Double-drift condition. Using the same conventions as in the plots in the main text, a positive angular difference indicates a shift toward the direction of the internal motion. Only trials where the speed of the gaze movement exceeded 2 standard deviations of the gaze speed in the pre-stimulus baseline are included. These distributions suggest that there was, at least on some trials, a pre-saccadic pursuit response broadly tuned to the direction of the target. However, these pre-saccadic responses in our dataset were too small and variable to discern whether the internal motion of the target in the double-drift condition biased the responses.

**Figure S4:**
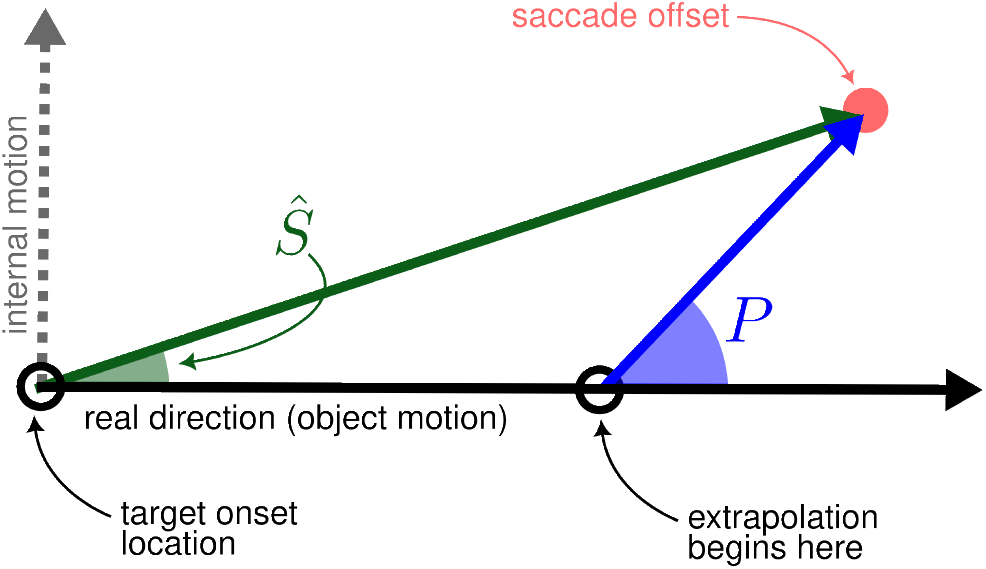
We estimated an extrapolation interval, corresponding to how far in time the saccadic system predicts future target locations. This analysis assumes that the saccade sistem extrapolates the target position using a biased motion equivalent to that driving early pursuit, and that this extrapolation begins only after a certain time has elapsed since the target onset. For each participant we found the interval *t* which minimizes the squared error between the observed direction bias, estimated by our analysis of saccadic landings, and the expected direction bias that would be obtained in the same analysis if the saccade extrapolated target position along the pursuit direction for an interval *t*^*\**^ (starting from an unbiased position). The expected saccadic direction bias Ŝ (shown as the green angle in the figure above) can be expressed as a function of the interval *t* and the pursuit direction bias *P* (shown as the blue angle in the figure above), 

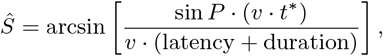

where *v* is the physical or external speed of the target, and latency and duration are the parameters of the saccadic eye movements. We found values of *t*^*\**^ numerically using data of the Experiment 2, separately for each participant; the mean value of *t*^*\**^ was 110 ms and the standard deviation across observers was 2 ms.

